# Genome Report: The reference genome of an endangered Asteraceae, *Deinandra increscens* subsp. *villosa*, endemic to the Central Coast of California

**DOI:** 10.1101/2024.02.25.582000

**Authors:** Susan L. McEvoy, Rachel S. Meyer, Kristen E. Hasenstab-Lehman, C. Matt Guilliams

## Abstract

We present a high-quality reference genome of the federally endangered Gaviota tarplant, *Deinandra increscens* subsp. *villosa* (Madiinae, Asteraceae), an annual herb endemic to the Central California coast. Stewards of remaining populations have planned to apply conservation strategies informed by whole genome approaches. Generating PacBio Hifi, Oxford Nanopore Technologies, and Dovetail Omni-C data, we assembled a genome of 1.67 Gbp as 28.7 K scaffolds with a scaffold N50 of 74.9 Mb. BUSCO completeness for the final assembly was 98.1% with 15.7% duplicate copies. We annotated repeat content in 74.8% of the genome. Long terminal repeats (LTR) covered 44.0% of the genome with *Copia* families predominant at 22.9% followed by *Gypsy* at 14.2%. Both *Gypsy* and *Copia* elements were common in ancestral peaks of LTR, and the most abundant element was a *Gypsy* element containing nested *Copia/Angela* sequenced similarity, reflecting a complex evolutionary history of repeat activity. Gene annotation produced 41,039 genes and 69,563 transcripts, of which >99% were functionally annotated. BUSCO duplication rates remained very high with proteins at 50.4% complete duplicates and 46.0% single copy. Whole genome duplication (WGD) synonymous mutation rates of Gaviota tarplant and sunflower (*Helianthus annuus*) shared peaks that correspond to the last Asteraceae polyploidization event and subsequent divergence from a common ancestor at ∼27 mya. Tandem genes were twice as prevalent as WGD genes suggesting tandem genes could be an important strategy of environmental adaptation in this species.

**Article Summary:** We introduce a high-quality reference genome for the endangered Gaviota tarplant. The assembly is 1.67 Gbp with 98.1% BUSCO completeness and 41 K annotated genes. We find extensive *Copia* long terminal repeat sequences and tandem genes that suggest environmental adaptation strategies. Comparisons with sunflower suggest a shared polyploidization event around 27 million years ago, close to the date of the common ancestor divergence. This work underlines the importance of genomic studies in accurately understanding adaptations and conservation needs.

## Introduction

Genetic analyses can play a critical role in the conservation of rare and endangered species (Theissinger et al. 2023). Regulatory agencies, land managers, and conservation biologists are increasingly turning to genetic analyses to inform decisions about the species they manage (Segelbacher et al. 2022). Management decisions take on new importance in light of the increasing threats to global biodiversity, such as the climate crisis (Scheffers et al. 2016), continued world-wide habitat destruction (González et al. 2020), and the spread of invasive species (North et al. 2021). Extending beyond traditional population genetic analyses, genome-scale data are being applied within a conservation framework. Genomic data results in more accurate characterization of genetic diversity and enables analyses dependent on genomic coordinates, functional annotations, and examination of features outside of the gene space, such as structural variants and regulatory elements (Fuentes-Pardo and Ruzzante 2017). Reference genomes combined with resequencing across a species’ range are being used, for example, to designate conservation units for the critically endangered *Artocarpus nanchuanensis* (Xia et al. 2023), to guide the selection of adaptive traits such as drought resistance in *Fagus sylvatica* (European beech; Pfenninger et al. 2021), to understand the adaptive potential in crop wild relatives (Wambugu and Henry 2022), and to aid in the restoration of locally-adapted *Castanea dentata* (American chestnut; Sandercock et al. 2022). The practice of conservation genomics will continue to increase as sequencing technology becomes more efficient and less expensive, computational resources and analytical techniques advance, and existing data and methodological resources grow via concerted efforts such as the Earth BioGenome Project (Lewin et al. 2018; Exposito-Alonso et al. 2020).

*Deinandra increscens* (H.M. Hall ex D.D. Keck) B.G. Baldwin subsp. *villosa* (Tanowitz) B.G. Baldwin (Madiinae, Asteraceae), commonly referred to as Gaviota tarplant, is an annual herb (Fig. 1) endemic to a narrow band along the coast in Central California, USA (Fig. 2). With only 25 total element occurrences, the taxon was listed under the California State Endangered Species Act (CESA; California Fish and Game Code sections 2050–2089.25) as Endangered in 1990 and the United States Federal Endangered Species Act as Endangered in 2000 (ESA; U.S. Code, Title 16, sections 1531–1544). According to the California Native Plant Society (California Native Plant Society 2023), the top threats to the taxon include development (11 EOs, or 44%), non-native plant impacts (6 EOs, or 24%), and road/trail construction and maintenance (5 EOs, or 20%).

**Figure 1.**
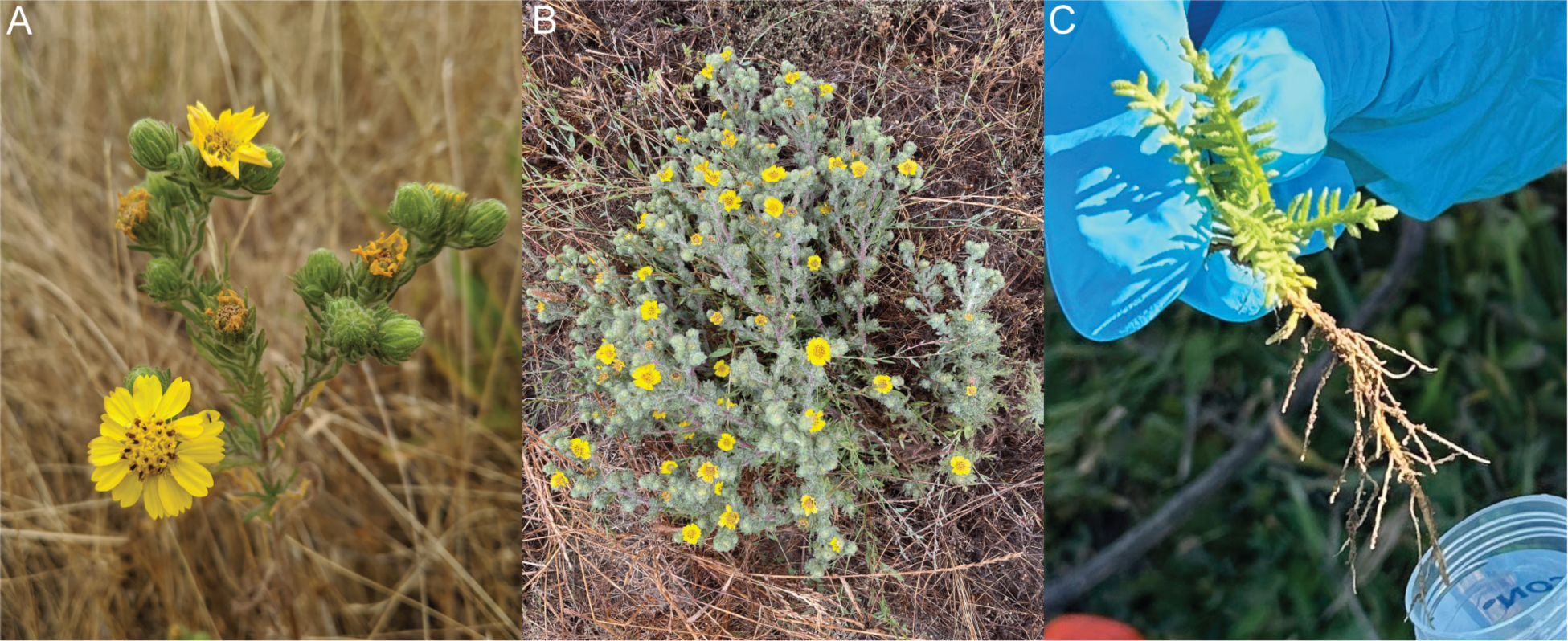
Examples of *D. increscens* subsp. *villosa,* which as an annual, can vary in form. A) Close-up of the inflorescence on a mature but smaller plant habit, B) An example of the larger plant habit. C) Seedling shoot and roots collected for RNA-Seq.

**Figure 2.**
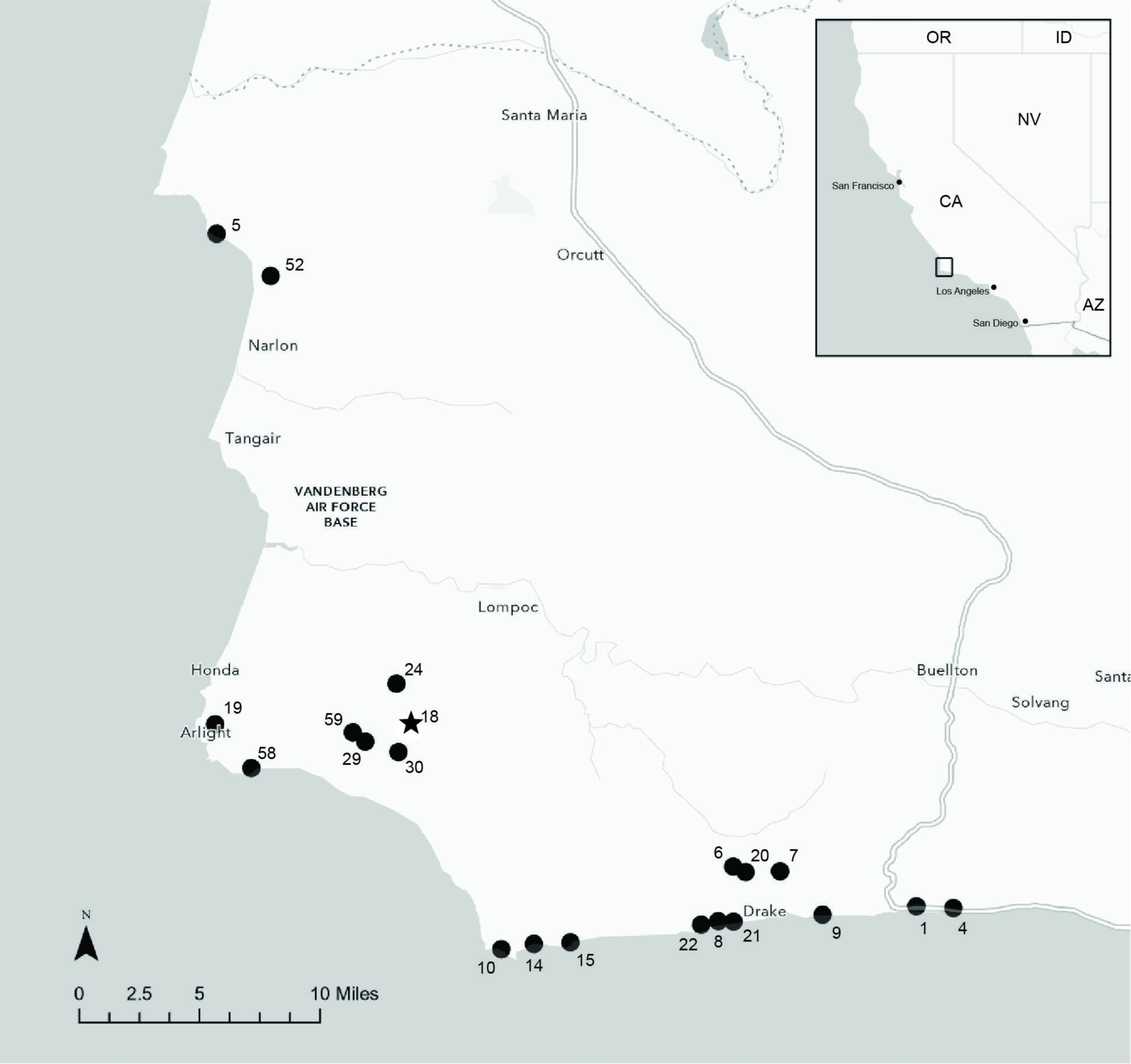
Map of southwestern Santa Barbara County, California, showing *Deinandra increscens* subsp. *villosa* element occurrences (EOs) as filled black circles, labeled with EO number. The star symbol indicates the EO where the reference individual was collected.

Molecular genetic approaches have long been applied to study members of subtribe Madiinae (including *Deinandra*), but to date, published studies have not included genome-scale data. The systematics and evolution of subtribe Madiinae have been extensively studied using phylogenetic analyses of Sanger sequence datasets (e.g., Baldwin 1992; Baldwin 1993; Baldwin and Wessa 2000; Baldwin 2005; Baldwin 2007), or less commonly using microsatellites (e.g., McGlaughlin and Friar 2011). Datasets using high-throughput sequencing are in development, however. Baldwin and colleagues report the development of a dataset of 100s of nuclear loci using a high-throughput, targeted sequencing approach to examine adaptive radiation of the tribe Madieae, inclusive of subtribe Madiinae (Baldwin et al. 2021). A large population genomics project by a subgroup of the authors is underway, using a genome resequencing approach to assist in conserving *D. increscens* subsp. *villosa* (McEvoy et al. 2023). Here we present the first step in that project, the development of a high-quality reference genome for *D. increscens* subsp. *villosa*.

## Methods & Materials

### Sample collection, preparation and sequencing

Fresh leaf and stem tissue were removed in the field from one mature individual of *D. increscens* subsp. *villosa*, vouchered on 7 May 2021 by Hasenstab-Lehman and Hazelquist (3314; SBBG) from the Tranquillon Mountain/Sudden Peak occurrence, south of Lompoc, CA, USA (34.575360, -120.520118, 416 m). Tissue was sampled into aluminum foil and immediately placed into a liquid nitrogen charged dry shipper for transportation to the Santa Barbara Botanic Garden (SBBG), where it was stored at -80F. The leaf tissue from this sample was used to generate all DNA sequencing data.

### Pacific Biosciences HiFi

High molecular weight (HMW) genomic DNA (gDNA) was extracted from a whole plant (1600 mg) using the method described in Inglis et al. (2018) with modifications. Sodium metabisulfite (1% w/v) was used instead of 2-Mercaptoethanol (1% v/v) in the sorbitol wash buffer and the CTAB buffer. Frozen tissue was ground in liquid nitrogen with mortar and pestle for 15 minutes, transferred to a 50 ml tube, and immediately mixed with 10 ml of sorbitol wash buffer. The suspension was centrifuged at 5000 x g for 5 minutes at room temperature and the supernatant was discarded. Using a paintbrush, the ground tissue pellet was gently resuspended in 10 ml of sorbitol wash buffer and this wash step was repeated 7 times to remove potential contaminants that may co-precipitate with DNA. The CTAB lysis step was performed at 45°C instead of 65°C for 1 hour with gentle inversion every 15 mins. The final DNA yield (17.3 µg) was quantified using a Quantus Fluorometer (QuantiFluor ONE dsDNA Dye assay; Promega, Madison, WI). The size distribution of the HMW DNA was estimated using the Femto Pulse system (Agilent, Santa Clara, CA) and 55% of the DNA fragments were found to be 30 Kb or longer.

The PacBio HiFi data generation was performed at UC Davis DNA Technologies Core (Davis, CA). The HiFi SMRTbell library was constructed using the SMRTbell Express Template Prep Kit v3.0 (Pacific Biosciences, Menlo Park, CA; Cat. #102-182-700) according to the manufacturer’s instructions. HMW gDNA was sheared to 15-18 Kb using Diagenode’s Megaruptor 3 system (Diagenode, Belgium; cat. B06010003). The sheared gDNA was concentrated using 1x of SMRTbell cleanup beads for the repair and a-tailing incubation at 37°C for 30 minutes and 65°C for 5 minutes. This was followed by ligation of overhang adapters at 20°C for 30 minutes, a second clean-up using 1x beads, and nuclease treatment at 37°C for 15 minutes. The SMRTbell library was size selected using 3.1x of 35% v/v diluted AMPure PB beads (Pacific Biosciences, Menlo Park, CA; Cat. #100-265-900) to progressively remove SMRTbell templates <5 Kb. The resulting library was sequenced using three 8M SMRT cells (Pacific Biosciences, Menlo Park, CA; Cat #101-389-001), Sequel II sequencing chemistry 2.0, and 30-hour movies on a PacBio Sequel II sequencer.

### Oxford Nanopore Technologies (ONT)

High molecular weight gDNA was extracted from the same reference individual as described above for HiFi. For ONT, 700 mg of frozen tissue was used, the suspension was centrifuged at 3500 x g for 5 minutes, and the wash step was repeated three times. The extracted HMW gDNA was purified using high-salt-phenol-chloroform (PacBio) to remove coprecipitating contaminants. Purity of the gDNA was checked using NanoDrop ND-1000 spectrophotometer and a 260/280 ratio of 1.89 and 260/230 ratio of 1.90 was observed. Total DNA yield was 2.5 µg as quantified by Qubit 2.0 Fluorometer (Thermo Fisher Scientific, MA). The integrity of the gDNA was verified on a Femto pulse system (Agilent Technologies, Santa Clara, CA) where an average fragment size of 11 Kb was observed.

The ONT sequencing library was prepared from 1.5µg of high molecular weight gDNA using the ligation sequencing kit SQK-LSK114 (Oxford Nanopore Technologies, Oxford, UK) following the manufacturer’s instructions with the exception of extended incubation times for DNA damage repair, end repair, ligation and bead elutions. 30 fmol of the final library was loaded on the PromethION R10.4.1 flow cell (Oxford Nanopore Technologies, Oxford, UK) and the run was set up on a PromethION P24 device using MinKNOW 22.10.7. Data was collected for 72 hours and basecalling was performed during the run using ONT’s basecaller ont-guppy-for-promethion 6.3.9.

### Dovetail Omni-C

Tissue from the reference sample was prepared using the DovetailTM Omni-C^TM^ Kit (Dovetail Genomics, Scotts Valley, CA). The manufacturer’s protocol was followed with slight modifications. Tissue was ground in liquid nitrogen with a mortar and pestle. Nuclear isolation followed published methods (Workman et al. 2019). Chromatin was fixed in the nucleus and digested under various conditions of DNase I until a suitable fragment length distribution of DNA molecules was observed. Chromatin ends were repaired, ligated to a biotinylated bridge adapter, and followed by proximity ligation. Crosslinks were reversed and the DNA was purified from proteins and treated to remove biotin external to ligated fragments. The library was generated using an NEB Ultra II DNA Library Prep kit (NEB, Ipswich, MA) with an Illumina-compatible y-adaptor. Fragments containing biotin were then captured using streptavidin beads, and the post-capture product was split into two dual-unique indexed replicates before PCR enrichment to preserve library complexity. The library was sequenced at the UCLA Technology Center for Genomics and Bioinformatics (TCGB; Los Angeles, CA) on an Illumina NovaSeq SP at 2X150, aiming to generate 400 million reads.

### RNA-Seq

Samples were collected from three individuals in the Tranquillon Mountain/Sudden Peak occurrence south of Lompoc, CA, USA (GVTP_1, GVTP_2 at 34.580914, -120.514450 and GVTP_5, 34.573121, -120.522042). Shoot and root tissue samples were collected from seedlings in rosette form and immersed in liquid nitrogen followed by -80 storage. In addition, leaf and stem tissues were collected from the adult reference individual already in freezer storage bringing the total number of samples to eight. Processing and sequencing was conducted by the UCLA Technology Center for Genomics and Bioinformatics. RNA was isolated from the frozen plant tissues using the Qiagen RNeasy Plant Mini Kit (Ref #74903) using lysis buffer RLT and 2M dithiothreitol (DTT). Library preparation of each sample was conducted using the KAPA Stranded mRNA-Seq Kit (96rxn; cat # KK8421). Sequencing of individual samples was generated at TCGB on an Illumina NovaSeq SP at 2X150 with two libraries per lane, aiming for 20 M reads per library.

### Genome assembly

Genome size estimation was conducted using both cytometric and sequencing methods. Flow cytometry trials were conducted comparing leaf and petal tissue of *D. increscens* subsp. *villosa*. Petals produced a better peak and were run using *Helianthus annuus* (sunflower) petal tissue and *Zea mays* (corn) and *Lactuca sativa* (lettuce) leaves as reference points. Resulting genome size estimates were used to predict the amounts of sequence data necessary for genome assembly. HiFi, and later ONT data, were used for sequence-based genome size estimation and ploidy using GenomeScope2 and Smudgeplot v0.2.2 (Ranallo-Benavidez et al. 2020). K-mer histograms of various sizes were created with KMC 3 (Kokot et al. 2017) to compare results. Contigs assembly trials were conducted initially with HiFi using the assemblers HiFiasm 0.16.1-r375 (Cheng et al. 2021), Flye v2.9.1-b1780 (Kolmogorov 2019), and wtdbg2 v2.5 (Ruan and Li 2020). To improve results, ONT sequencing was generated and run in Flye and Canu v2.2 (Koren et al. 2017) in combination with HiFi reads in Verkko v1.4.1 (Rautiainen et al. 2023). ONT-based Flye, Canu, and ONT/HiFi Verkko assemblies were run through Purge Haplotigs v1.1.2 (Roach et al. 2018) using Minimap2 v2.24-r1122 (Li 2018) alignments to separate out under-collapsed haplotype duplication. Of these, the Flye and Canu resulted in the best contig assemblies and were tested in the scaffolding step.

Scaffolding followed recommended methods and tools for the 3D-DNA pipeline (https://aidenlab.org/assembly/manual_180322.pdf). Omni-C reads were aligned to the contig assembly with BWA (Li 2013) and these were used as input for Juicer v1.5 (Durand et al. 2016) to create a formatted file of duplicate-filtered long-distance pairs. This pairs file was provided along with the genome to 3D-DNA v180922 for scaffolding (Dudchenko et al. 2017). Tuning of parameters was evaluated by visualization of Omni-C alignment heatmaps in JuiceBox (Dudchenko et al. 2018). Insufficient Omni-C data over greater portions of the Canu contig assembly was the rationale for eliminating this version and the Flye assembly was selected for the remaining steps.

The scaffolded ONT-based Flye assembly was screened for assembled chloroplast using both chloroplast contigs assembled by GetOrganelle v1.7.7.0 (Jin et al. 2020) and the closest relative with a chloroplast reference genome, Asteraceae *Achyrachaeana mollis* (NCBI RefSeq accession NC_036504.1; Persinger et al. 2017). Contaminant screening was done with both Blobtools2 v4.1.3 and the NCBI Foreign Contamination Screening tools FCS-GX v0.4.0 (Astashyn et al. 2023). Gap filling was done using TGS-GapCloser v1.2.1 (Xu et al. 2020).

Throughout the workflow, contig and scaffold assemblies were evaluated with QUAST v5.2.0 (Gurevich et al. 2013) and BUSCO v5.4.3 using the embryophyta_odb10 database (Manni et al. 2021). Final assembly completeness was also evaluated with Merqury v1.3 (Rhie et al. 2020). This involved removing the bridge adapters from the Omni-C reads using Cutadapt v2.6 (Martin 2011) to prepare histograms of k-mer counts in Meryl v.1.3 (Rhie et al. 2020). A full list of software and versions used in the genome assembly process is also available in Supplemental Table 1.

### Repeat annotations

A consensus of transposable elements (TEs) and other repetitive content was identified *de novo* throughout the final reference genome using RepeatModeler v2.0.4 (Flynn et al. 2020). These repeats were combined with curated plant long terminal repeats (LTRs) in InpactorDB non-redundant v5 (Orozco-Arias et al. 2021) and softmasked with RepeatMasker v4.1.5 (Smit et al. 2013). The RepeatMasker .out format file was used as input for the script ParseRM (Kapusta 2017) to generate summary statistics according to TE category and to format data for the TE landscape abundance, which was plotted in ggplot2 v3.4.2 in R v4.2.1 (Wickham 2016).

### Gene annotations

RNA-Seq alignments were used to improve gene prediction. RNA-Seq reads were checked for quality control with FastQC v0.11.9 (Andrews 2010) and then trimmed with fastp v0.22.0 (Chen et al. 2018). Initial alignments with Hisat2 v2.2.1 (Kim et al. 2019) were lower than the desirable threshold of 80%, so the unmapped reads were screened for contaminants with Kraken2 v2.1.2 (Wood et al. 2019) using the MiniKraken v2 database of bacteria, archaea, and virus. Matches were removed from the subset of reads that did not map using seqtk v1.3-r106 (Li 2023) and the remaining reads were combined back with the mapped readset. These improved read alignments were provided as input along with the soft-masked genome for the EASEL v1.3 annotation pipeline (Webster et al. 2023) using the default recommended configuration for plants with the exception of a reduced “rate” value of 70 to include all aligned libraries. EASEL assembles reference-based transcripts using Stringtie2 (Shumate et al. 2022) and PsiCLASS (Song et al. 2019), identifies open reading frames with TransDecoder (Haas 2023) refined by EggNOG-mapper (Cantalapiedra et al. 2021), and uses resulting transcript and protein alignments for training and gene prediction with AUGUSTUS (Stanke et al. 2006) resulting in an unfiltered structural annotation with alternative transcripts. A matrix of primary and secondary sequence features for resulting transcripts is scored and filtered using a random forest algorithm with a plant-based training set. Final metrics are generated using AGAT (Dainat et al. 2022), gene space completeness with BUSCO, and functional annotation with EnTAP (Hart et al. 2020).

### Gene duplication

To examine the potentially extensive gene duplication reflected in the BUSCO score of the final assembly, genome-wide gene duplication was identified and categorized using DupGen-finder (Qiao et al. 2019). Genes categorized as whole genome duplication (WGD) were used to calculate and plot the frequency of duplication by synonymous mutation (Ks) to identify potential WGD events. WGD paralogs were also visualized in Circos (Krzywinski et al. 2009) to examine their genome-wide distribution.

## Results and Discussion

### Sequencing and genome assembly

Three HiFi sequencing runs resulted in a total of 6.3 M reads with an N50 of ∼13 K, consisting of 76.8 B bases for 46x in coverage using the assembly length and ONT-based genome size estimates. Adaptor filtering only removed one read. ONT sequencing resulted in 14.8 M reads with an N50 of 10.7 K, 101.2 B bases, and a coverage of 61x. Two Omni-C lanes generated 848 M 150 bp PE reads in total with 126.8 B bases and 75.9x coverage.

Flow cytometry using related species of *L. sativa*, *H. annuus*, and *Z. mays* as general reference points and petal tissue of *D. increscens* subsp. *villosa* helped place it at an average 1C of 2.2±0.1 pg, which translates to a base count of approximately 1.7 to 2.3 Gb (File S2). This estimate was supported by sequence-based k-mer profiles where genome lengths in ONT trials using k-mers 21 and 35 were 1.61 Gb and 1.69 Gb, and HiFi trial with k-mers 21, 27, and 31 ranged from 1.77 Gb, 1.76 Gb to 1.74 Gb (Fig. 3a). Smudgeplots created with the same histograms all supported a diploid genome (Fig. 3b).

**Figure 3.**
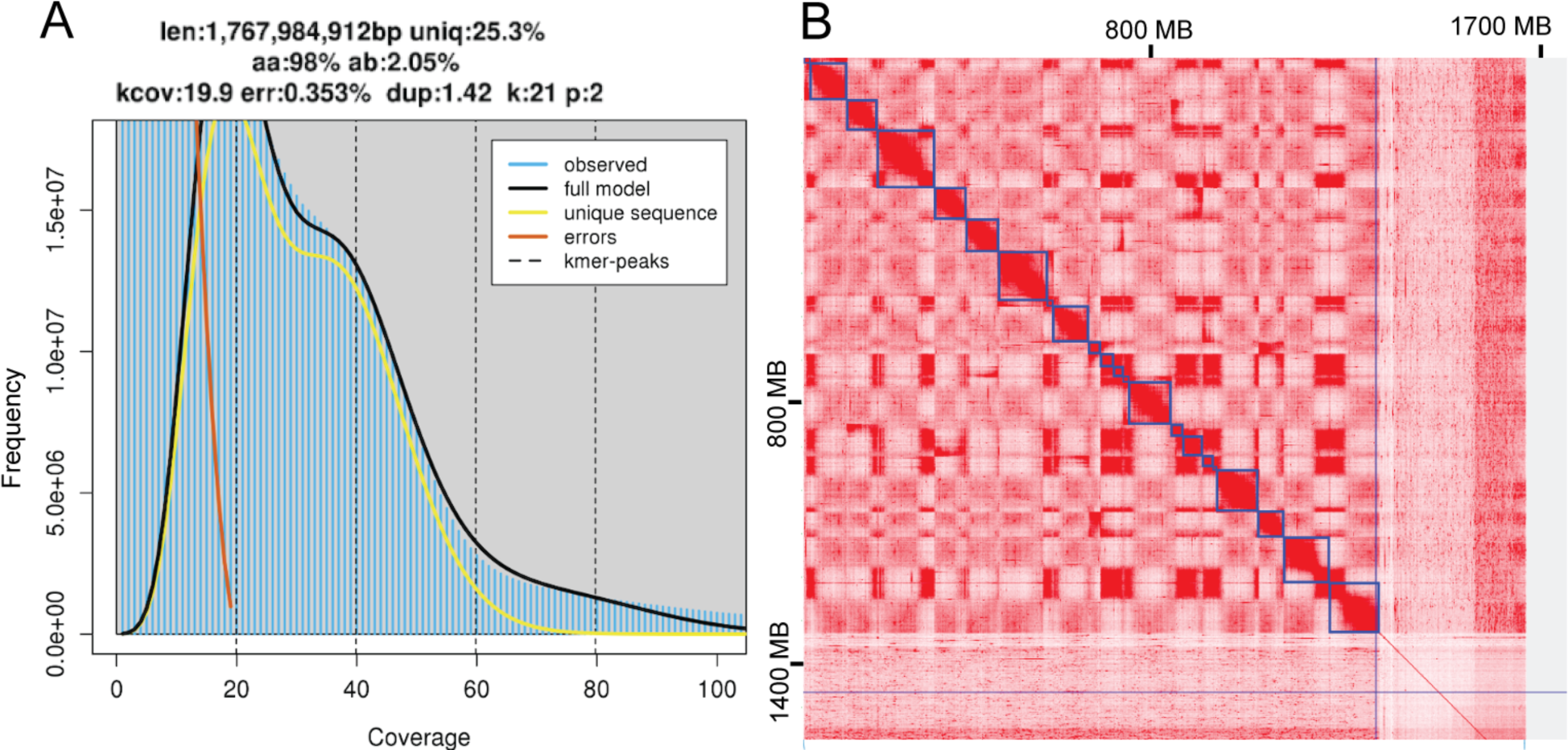
A) Genome estimates generated with HiFi sequencing using the diploid setting with k-mer size 21. Note the double peaks with the top of the first peak cut off and overlapped by the error model (red line). Genome size was estimated at 1.77 Gbp with 75% repeat content and 2% heterozygosity. B) Heatmap of Omni-C read pairs mapped to the scaffolded genome. The X-axis shows a total length of ∼1650 Mbp. Blue lines delineate the 21 largest scaffolds estimated by 3D-DNA. Discarded fragments and small contigs to the right show a thin line of alignment indicating an absence of significant Omni-C support for anchoring in superscaffolds.

The contig assembly created with Flye was 2.46 Gbp in length with 101 K contigs and an N50 of 76.6 Kb. While fragmented, the BUSCO embryophta score was 99.2% complete, 79.2% of which was duplicates. Purging undercollapsed haplotypes reduced the BUSCO score to 98.3% with 22.2% duplicates. The purged assembly was 1.67 Gbp in 37 K contigs with an N50 of 108 Kb. The alternate haplotype assembly was only 75% complete and was not processed further. The combined primary and alternate haplotype assemblies resulted in a 71.48% completeness score in Merqury, with a QV of 28.79. With minor adjustments to remove contaminants, plastid contigs, and fill gaps, the primary scaffolded assembly was 1.67 Gbp as 28.7 K scaffolds with a scaffold N50 of 74.9 Mb, L50 of 9, and 600 Ns per 100 Kbp (Table 1). Statistics for other assembly trials are available in File S1. BUSCO completeness for the final assembly was 98.1% with 15.7% duplicate copies, 0.9% fragmented, and 1.0% missing. Assessment of the primary haplotype assembly in Merqury was 62.25% with a QV of 29.49, likely reflecting similar issues as seen with the other k-mer based analyses like GenomeScope.

**Table 1.**
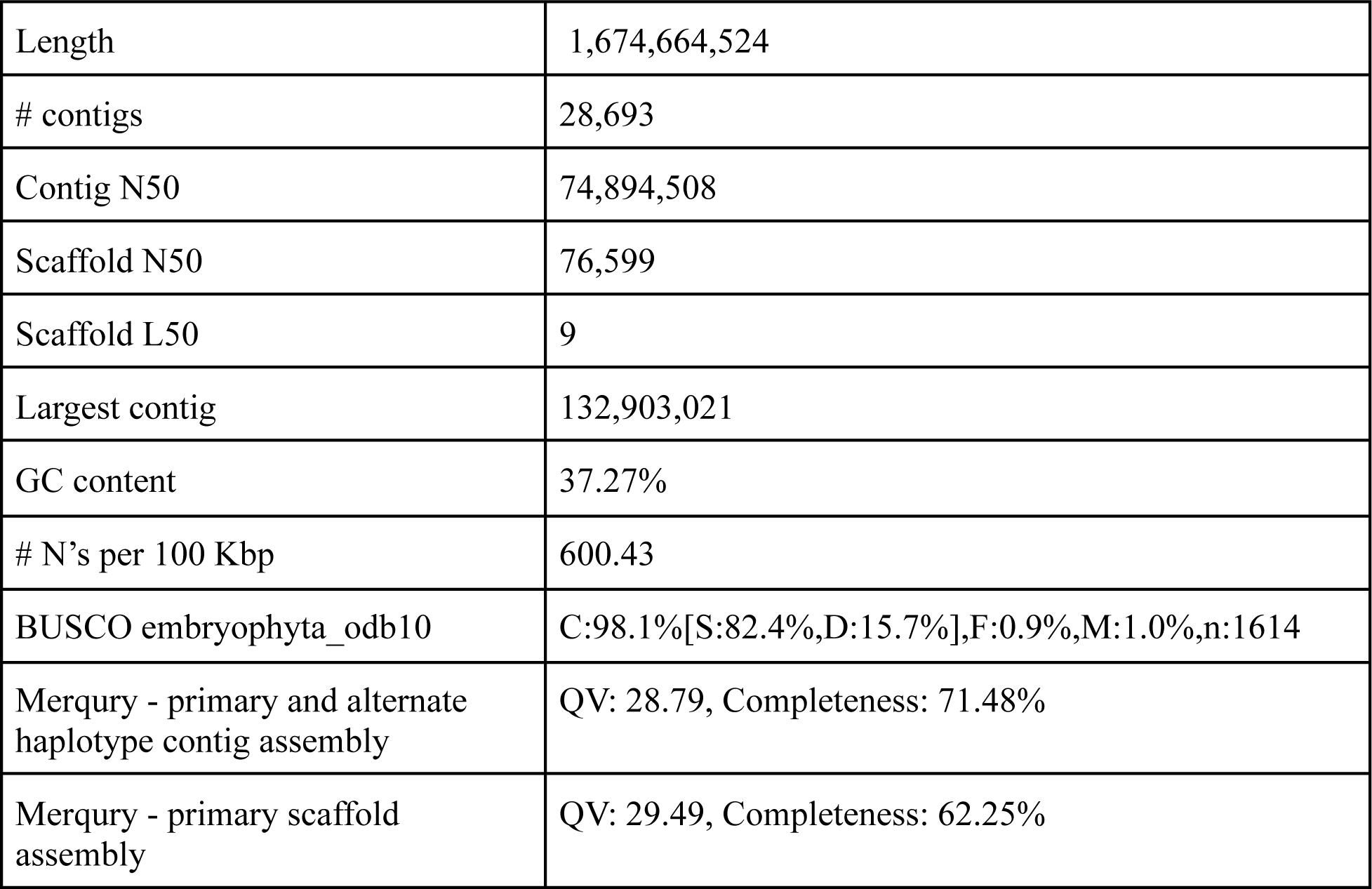
Genome assembly statistics.

### Repeat annotations

Repeats were identified in all but 53 scaffolds and softmasked in 74.81% of the genome (Table 2). LTRs covered 43.96% of the genome with *Copia* families predominant at 22.88% followed by *Gypsy* at 14.15% (Fig. 4a). There were more copies of *Copia* overall, but fewer unique, with 451,119 copies identified within 29,511 unique elements. *Gypsy* had 272,599 copies total with 32,255 unique elements. DNA transposons masked 2.6%, led by MULE-MuDr (0.76%), CMC-EnSpm (0.59%), and PiF-Harbinger (0.56%). A TE landscape abundance plot, using divergence to represent relative time of insertion, revealed more ancestral activity compared to recent (Fig. 4b).

**Table 2.**
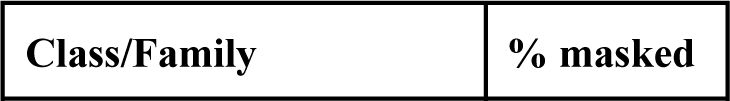
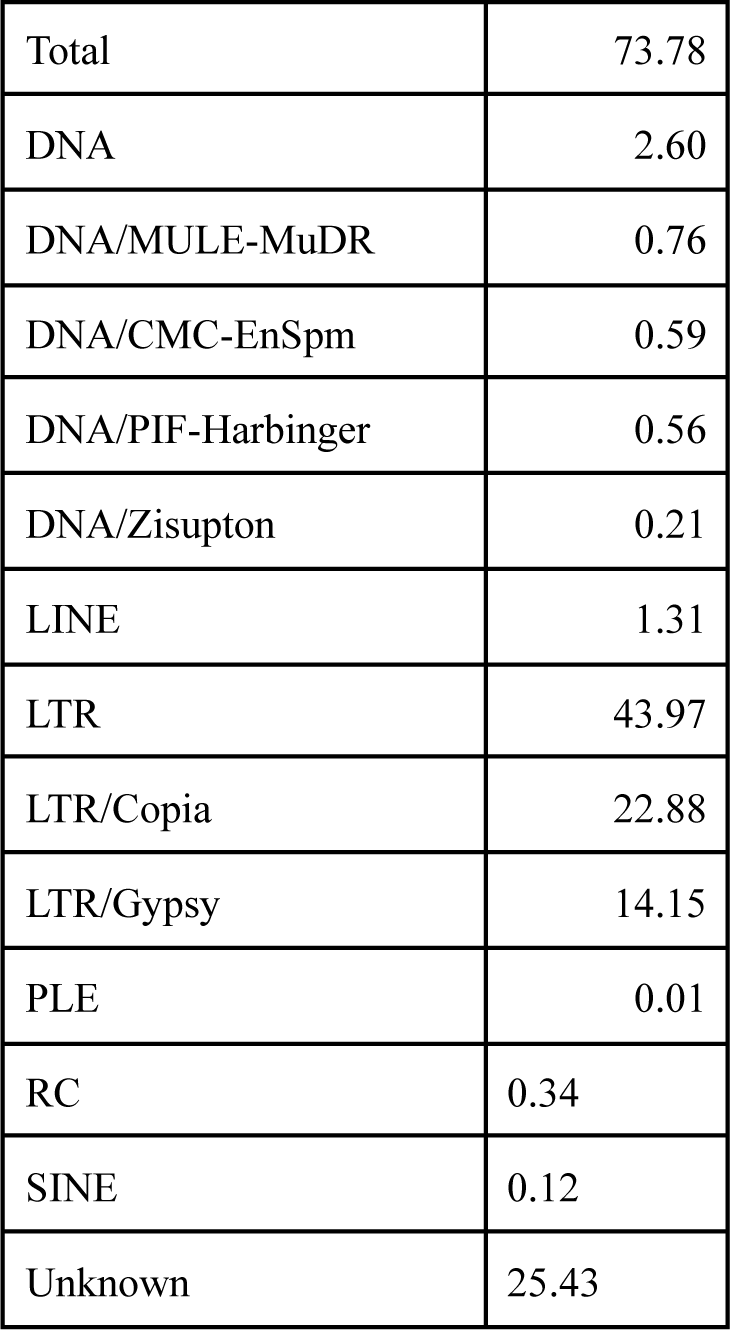
Repeat annotation statistics for all TE classes and most abundant families.

**Figure 4.**
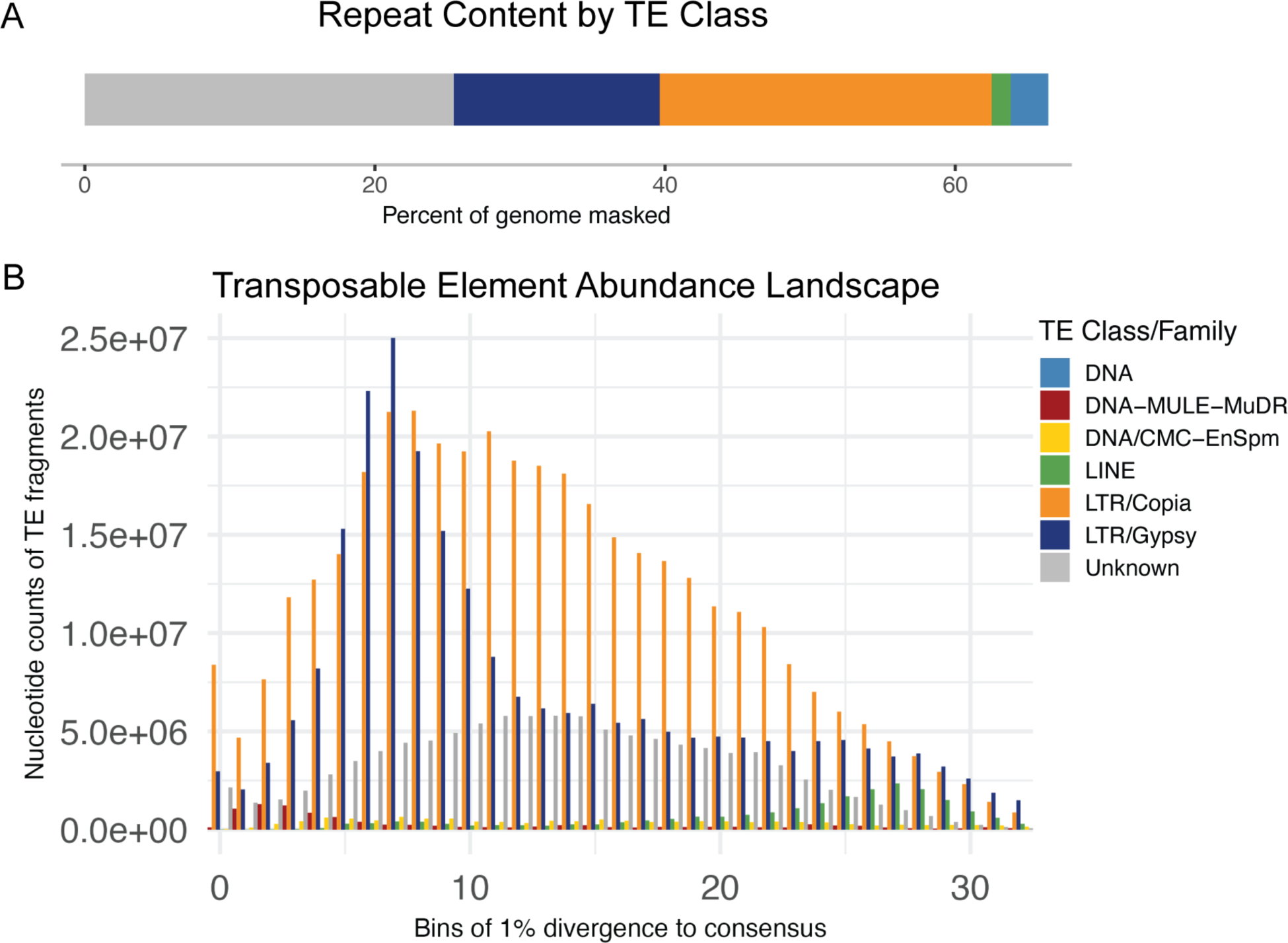
A) Percentage of base pairs masked as transposable elements (TE) across the *D. increscens* subsp. *villosa* genome. TE classes shown: Unknown, LTR/*Gypsy* LTR/*Copia*, LINE, and DNA, 74.8% in total. B) Abundance of TE elements as base pair totals binned by divergence as a proxy for relative time. Only the most abundant TE classes and families are plotted in this figure; the full summary of elements can be found in File S1.

Interestingly, elements found most frequently within these ancestral peaks of abundance were from a mix of classes and families including both LTRs *Gypsy/Tekay* and *Copia/Sire*. Two of the most prevalent elements in these peaks were unclassified with only small sections of sequence similarity to anything within ASTER-REP, a curated Asteraceae-specific TE database created from genome skimming (Ventimiglia et al. 2023). Recently active TEs were also a mix led by *Copia/Angela*, followed by *Gypsy/Tekay* and unclassified elements. *Copia/Angela* elements were more accurately identified by sequence similarity using *Helianthus* elements in the ASTER-REP database (Ventimiglia 2022). The element with the greatest peak of abundance was originally identified as *Gypsy* by RepeatMasker, but was also found by sequence similarity to have an internal *Copia/Angela* element. Such complex, nested, or unknown elements are likely indicative of the long history and high level of repeat content found within the lineage.

The amount and composition of TE content in a genome are due to the success of different TEs active within that genome. While genomic studies do not exist for most Asteraceae, comparisons of transposable elements exist for important crop species including *L. sativa*, *Cynara cardunculus* var. *scolymus* (artichoke), *Artemisia annua*, (sweet wormwood), *Carthamus tinctorius* (safflower), and *Chrysanthemum seticuspe* (Ventimiglia et al. 2023). In particular, much work exists on the more closely related *H. annuus* (Badouin et al. 2017; Mascagni et al. 2017; Ventimiglia et al. 2023) which has a divergence time of 26.9 mya from *D. increscens* (Kumar et al. 2017). *H. annuus* and *D. increscens* subsp. *villosa* are both at the high end of total repeat content for Asteraceae, at almost 80%. They are also similar in LTR proportion, at ∼42%, which falls mid-range for these Asteraceae (35.27% in *C. cardunculus* var. *scolymus* and 52% in *C. seticuspe*). But ratios of *Gypsy* to *Copia* vary dramatically across Asteraceae with some favoring one family over the other. *H. annuus* is on one end of the Asteraceae spectrum, with a ratio of 6.47 *Gypsy:Copia* (only 6.35% *Copia*), and *D. increscens* subsp. *villosa* on the other with 0.62, which further supports the hypothesis that different LTR sublineages had different rates of activity after speciation (Ventimiglia et al. 2023). TE activity has greatly influenced genome evolution in both these species with very different results.

### Gene annotations

After contaminant filtering, read counts of the eight RNA-Seq libraries ranged from 81M to 130M with alignment rates ranging from 76.0% to 91.42%. From this, StringTie generated 76,471 transcripts while PsiCLASS produced 116,777. Before filtering, the consensus of transcripts totaled 334 K with 197 K genes, containing many false positives as indicated by the high mono to multi-exonic rate (0.69) (Vuruputoor et al. 2023) and low sequence similarity rate within EnTAP (0.65). False positives were well resolved by filtering, with only 69,563 final transcripts and 41,039 genes, with a more typical mono/multi-exonic rate of 0.24 and a high 0.92 sequence similarity rate reported by EnTAP (Table 3). Over 99% of the transcripts had functional annotations provided by sequence similarity and/or gene family assignment. Even after filtering, BUSCO duplication rates remained very high with 71.8% duplication and 24.8% single copy among transcripts (96.6% total) and proteins at 50.4% complete duplicates and 46.0% single copy (96.4% total complete). Despite the large number of assembly scaffolds, genes are only present in 2,444 scaffolds and 92% of the total genes are consolidated in the 22 largest. Previous cytogenic chromosome counts including one sample of *D. increscens* subsp. *villosa* and multiple samples of subsp. *increscens* to be 1N = 12 (Tanowitz 1982), so these 22 contigs may represent the significant portions of pseudo-chromosomes.

**Table 3.** Gene annotation statistics.

|  |  |
| --- | --- |
| Filtered: Total number of genes: | 41039 |
| Total transcripts | 69563 |
| EnTAP alignment rate: | 0.92 |
| Mono-exonic/multi-exonic rate: | 0.24 |
| BUSCO (embryophyta_odb10): | C:96.6%[S:24.8%,D:71.8%],F:0.4%,M:3.0%,n:1614 |

### Gene duplication

Protein-coding gene pairs were grouped into the following categories: 4,608 WGD pairs, 9,879 tandem pairs, 2,353 proximal pairs, 5,368 transposed pairs, and 30,457 dispersed pairs. Plotting links between the WGD gene pairs revealed that they are mostly, but not exclusively, found in gene-dense regions on the 22 largest scaffolds plus two others that are very short (Fig. 5A). The WGD pairs produced two peaks of elevated frequency at synonymous mutation rates (Ks) 0.66 and 1.60 that aligned approximately with peaks in *H. annuus* at 0.59 and 1.81 (Fig. 5B). In *H. annuus*, Badouin et al. (2017) used paralogs and homologs from speciation events to identify the most ancient peak as the well-established core eudicot WGT-ɣ event. The more recent peak is two overlapping polyploidization events from the Asteraceae lineage: WGT-1 at 38-59 mya and WGD-2 at 29 mya. The most recent peak, observed only in *H. annuus* (the green line in Fig 5B), was reported as tandem genes. Our method removed non-WGD genes like tandems before plotting Ks values, so this peak is not portrayed, but tandem genes in *D. increscens* subsp. *villosa* are elevated even beyond WGD at over 15 K genes (9,879 pairs), or 38% of the total protein-coding genes. The frequency of tandem duplications closely matches gene-dense regions (Fig. 5A).

**Figure 5.**
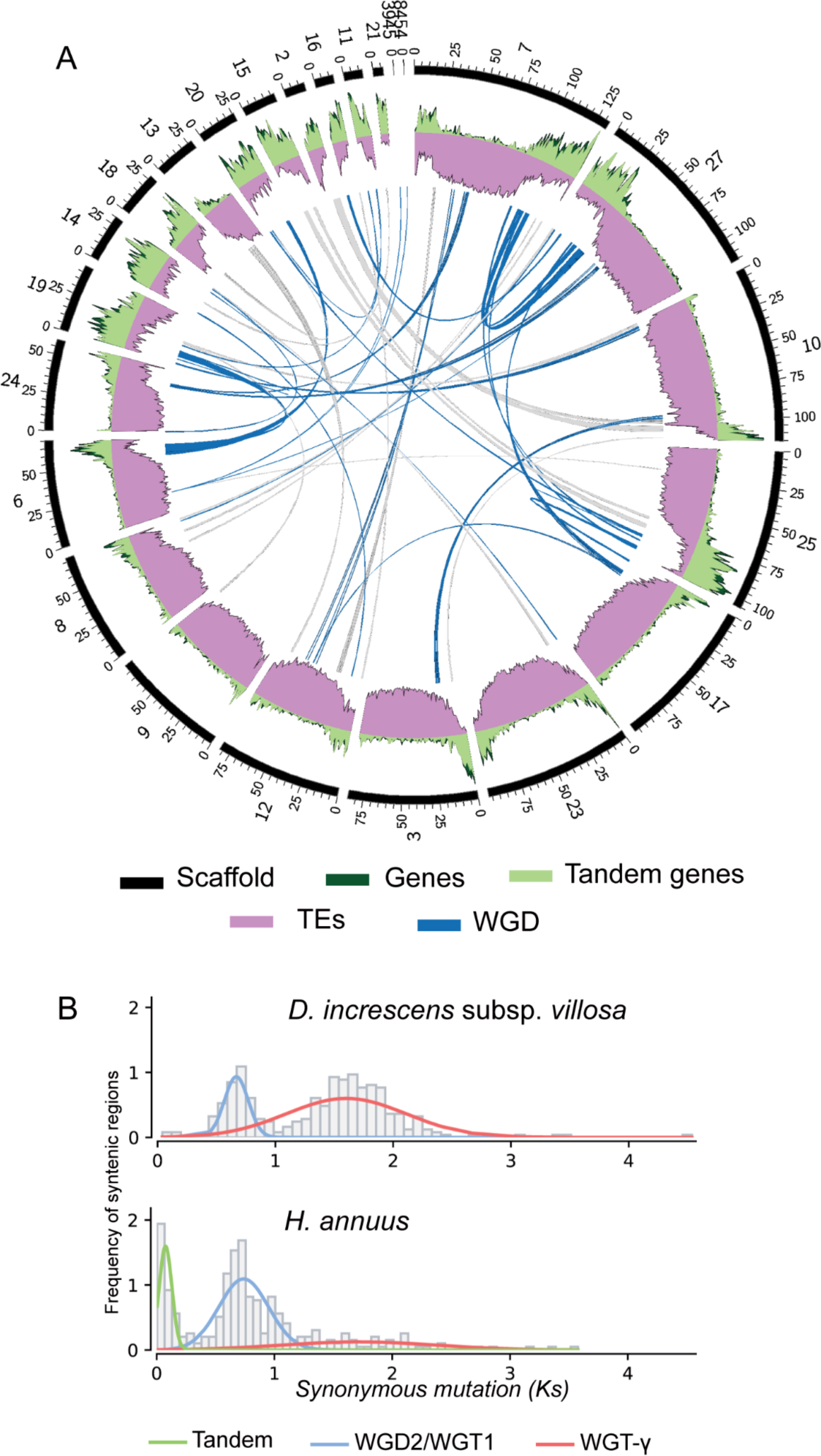
A) Circular plot representing the 22 largest scaffolds (black) clock-wise by size plus two others all containing whole genome duplication (WGD) gene pairs linked in blue. Gene density (counts per 100 Mbp window) is plotted in dark green and overlaid by the density of tandem genes in light green. Transposable element (TE) frequency as a percentage of base pairs per window is shown in purple. B) Peaks representing whole genome duplication events in *D. increscens* subsp. *villosa*, (top), and *H. annuu*s (bottom). Peaks consist of an increased frequency of syntenic blocks, which are regions of collinear genes in proximity, by the rate of observed average rate of synonymous mutations for the blocks.

WGD and tandem genes differ in their evolutionary consequences. In angiosperms, WGD events have been shown to be followed by diversification of species (Landis et al. 2018) which could in part explain the timing of the divergence of *H. annuus* and *D. increscens* lineages after the two more recent whole genome events. WGD genes and tandem genes can play significant and sometimes complementary roles in adaptation to environmental changes (Guo et al. 2022). They may also have functional bias, with tandem genes containing lineage-specific adaptations to environmental stimuli in areas of stress response and disease resistance as compared to the more regulatory roles of WGD genes as observed in *Populus trichocarpa* and *Arabidopsis thaliana* (Hanada et al. 2008; Rodgers-Melnick et al. 2012; Qiao et al. 2019).

### Summary

In this study, we leveraged long-read DNA sequencing and Hi-C data to generate a scaffold-level reference genome assembly for the highly repetitive and heterozygous *D. increscens* subsp. *villosa*. The reference includes repeat annotations and protein-coding gene annotations, along with high-level characterization of extensive gene duplication throughout the genome. Larger genomes with a high percentage of repeats can result in assemblies not achieving chromosome-level status as was seen in the related *H. annuus*, which has a similar total repeat content. While the haploid version of the *D. increscens* subsp. *villosa* genome presented here is highly fragmented, much of the fragmentation seems to exist within the repeat space and the majority of gene space is consolidated within the 22 largest scaffolds. Complexity of repeat content and heterozygosity could be further resolved with ultra-long read sequencing. Additional sequencing coverage, or chromosome counts, could also clarify conflicting evidence of ploidy level. While the total repeat content is similar to *H. annuus*, TE lineage abundances are dramatically different, indicating activity of a variety of TEs classes and families throughout time with very different outcomes. Tandem and whole genome duplication is also described within the context of *H. annuus* and other Asteraceae, revealing evidence of the WGD and WGT events shared by this family and likely occurring only a few million years before divergence of the *D. increscens* subsp. *villosa* lineage. The abundance of tandem gene duplications is particularly interesting and could be a source of defense response and other environmental stress response related genes. Whole genome annotation of genes, repeats, and gene duplications, especially the high number of tandems will provide a valuable reference for future analyses, and more accurate results for current efforts involving resequencing the broader element occurrences for genetic and genomic insights to inform conservation monitoring and mitigation.

## Supporting information

File S1 - Workflow output summaries

## Data Availability

Data is available under NCBI BioProject ID PRJNA1046654 with reference genome reads in BioSample SAMN38504800 and additional RNA-Seq in BioSamples SAMN39983642-7. The reference assembly with gene annotations is available at https://github.com/slmcevoy/gaviota-tarplant/blob/main/manuscript/gitupload.tar.gz (NCBI accession TBD). Methods and commands are at https://doi.org/10.5281/zenodo.10702999. Supplemental material is at https://github.com/slmcevoy/gaviota-tarplant/blob/main/FileS1-summaries.xlsx.

## Acknowledgments

We thank the United States Fish and Wildlife Service for support of the Santa Barbara Botanic Garden’s Endangered Species Act 10(a)1(A) recovery permit to work with *Deinandra increscens* subsp. *villosa* (Permit No. TE-094893-3). We similarly thank the California Department of Fish and Wildlife for support of the Garden’s Scientific, Educational, or Management Permit to work with State-listed rare plants (No. 2081(a)-16-015-RP). For assisting with access in the field, we thank Jennifer Garcia, Elihu Gevirtz, and Richard Podolsky of BayWa r.e.; Kelly Schmoker of California Department of Fish and Wildlife; Jake Marcon of Dudek; Elizabeth Hiroyasu, Moses Katkowski, Chelsea Nielsen, and Laura Riege of The Nature Conservancy’s The Jack and Laura Dangermond Preserve; and Luanne Lum of Vandenberg Space Force Base. We thank Elijah Balderas, Susana Delgadillo, Emily Thomas, Caitlin Hazelquist, and Isabel Rivera for assistance with plant tissue collections in the field and Annie Ayers for help managing specimens in the herbarium at the Garden and creation of the element occurrence map. We are grateful to Bruce Baldwin, Mark Elvin, Barry Tanowitz, and Dieter Wilken for discussions about the taxonomy of *D. increscens* subsp. *villosa.* We thank Ohalo Genetics for conducting the flow cytometry analysis, especially Sandra Owusu. DNA long-read extraction, library preparation, and sequencing were carried out at the DNA Technologies and Expression Analysis Cores at the UC Davis Genome Center by Noravit Chumchim (HiFi library preparation), Oanh Nguyen (HiFi sequencing) and Ruta Sahasrabudhe (ONT), supported by NIH Shared Instrumentation Grant 1S10OD010786-01. Omni-C libraries were prepared by Will Seligman of the UCSC Paleogenomics Lab and sequenced by the UCLA Technology Center for Genomics & Bioinformatics who also handled generation of RNA-Seq. Assembly and annotation were carried out with the computational resource support provided by: UCSC Genomics Institute, Merly Escalona of the UCSC Paleogenomics Lab, and NSF ACCESS (Boerner et al. 2023) and Jetstream2 (Hancock et al. 2021) cyberinfrastructure.

